# Degree of urbanization and predation pressure on artificial lepidopteran caterpillars in Cali, Colombia

**DOI:** 10.1101/493924

**Authors:** Jefferson Cupitra-Rodriguez, Lorena Cruz-Bernate, James Montoya-Lerma

## Abstract

Growing urban expansion results in the alteration of ecological processes (i.e. predation) within trophic networks. Predation on herbivores is known to vary with the size of the area covered in vegetation, successional stage, altitude, and the structure of the predator community, but there are gaps in information regarding how this occurs in urban and suburban environments. The purpose of this study was to determine whether the predation pressure on artificial models of lepidopteran larvae varied with degree of urbanization, type of substrate, and group of predators (birds or arthropods) in Cali, Colombia. Five hundred and eighteen artificial larvae were placed in two areas of the city (urban vs. suburban) and in two types of substrate (leaf vs. stem) for 30 continuous days and with two replications over time. Total predation was measured as the number of models with evidence of attack by predators. The overall incidence of predation was 24.13%, and was significantly higher in the urban area (63.20%) when compared to the suburban area (36.80%). The leaf substrate was attacked significantly more than the stem (60% vs. 40%). The proportion of attacks carried out by birds was significantly higher (74.40%) than that carried out by arthropods (24.80%). Together, these results suggest that the incidence of predation varies with the disturbance caused by urbanization and by the type of substrate in which prey organisms are found. In addition, the study confirms that birds are the main controllers of herbivorous insects in urban environments.

## Introduction

The drastic transformation of pristine natural landscapes into fragments within urban landscapes [1–2] causes changes in community composition, structure, abundance, and trophic relations [3]. The size of these fragments of vegetation and the degree of disturbance in the interior and surrounding areas affect the permanence of species that require large continuous areas of habitats and particular food resources [4–6]. When species disappear from a habitat, predator-prey relationships change and some species are temporarily favored. For example, herbivore communities undergo rapid growth in habitats where populations of its natural enemies are low or have disappeared due to human induced changes [7, 9].

Predation pressure varies both within and between habitats due to differences in community and predator density, vegetation structure and complexity, and intensity of anthropogenic disturbance [2, 10–11]. Compared to the interior, the edges and patches of forest fragments are considered as areas of greatest predatory risk for many species [5, 8, 11, 12]. In forest clearings, herbivores and their predators are more abundant since these sites show increased leaf and plant growth compared to the understory [8]. When comparing forests with open and close canopies with rural, researchers [11] recorded that both nests and artificial caterpillars were attacked more with increasing level of disturbance. The degree of prey exposure also influenced detection by predators. For example, predatory rate on artificial larvae located exposed on leaves was greater compared to those that were hidden [13].

Usually predatory events happen quickly and are often hard to measure because predators may hide while consuming prey, or they may be nocturnal leading to lower detection. Similarly, the degree of predation is complex to measure because, frequently no traces or only fragments of the consumed prey are found. One technique to measure predator pressure is using dyed prey organisms that allows tracking of prey. However, one of the disadvantages of this method are the low number of individuals effectively tracked, time-intensive observation, and difficulty in identifying the natural enemies of prey [14]. To resolve these difficulties, we can use artificial plastic prey that conserve marks of attacks received by natural enemies on their surface. This method has been used to make estimates of predator pressure on caterpillars [8, 13–15]. The majority of these studies were carried out in wooded areas, with a different successional level, and few of them evaluated predatory pressure in urban and suburban environments [2,16]. It has been suggested that the calculation of predation rate on artificial caterpillars does not differ from that occurring in nature [8].

Establishing how urbanization affects incidence of predation provides knowledge regarding the dynamics of urban ecological interactions and offers a tool for the management of the populations involved. The main purpose of the present study was to determine whether predation pressure (estimated using artificial models) varies as an effect of anthropogenic disturbance. The study also sought to establish whether substrate type influences predation incidence and which type of predator, bird or arthropod, exerted greater predation pressure on artificial lepidopteran larva models.

## Materials and methods

### Study area

This study was carried out in Santiago de Cali, Department of Valle del Cauca, Colombia (3°32’33’’ N, 76°31’58’’ O; 995 masl) between August 2015 and August 2016. The city has a mean temperature of 24°C, average annual rainfall of between 1000 and 2000 mm, and a bimodal climate with dry periods during January-February and July-August and rainy periods during March-June and September-December. These climatic characteristics correspond to the dry tropical forest life zone (df-T) according to Holdridge [17].

### Predation tests

Two different areas were defined according to the degree of human disturbance: urban and suburban, and two sites were chosen in each one: Universidad del Valle Campus (UV) and Constructora Limonar (CL) in the urban area and, Parque de las Garzas (PG) and Hacienda Cañas Gordas (CG) in the suburban area. Ten “Chiminango” trees (*Pithecellobium dulce*) were selected at each site, with a distance of at least 30 m. between each tree. This species is one of the most common and most used by birds in the area [18–19]. Medium-sized artificial models (40 mm long and 6 mm in diameter) based on natural butterfly larvae, were made from odorless, non-toxic modelling clay (Fig 1).

**Fig 1.**
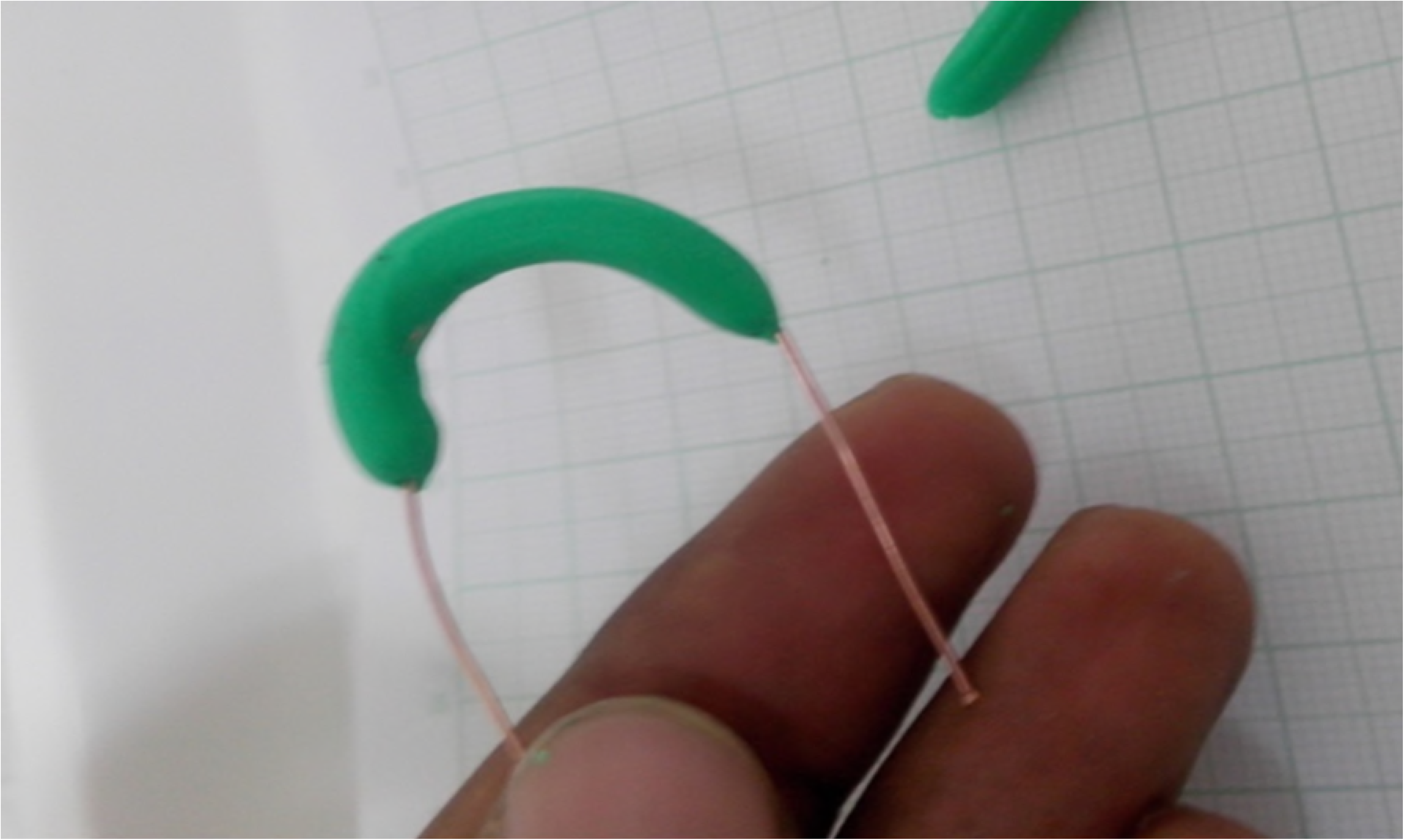
Artificial caterpillar model. Prototype (elongated) of the lepidopteran larva model used in the predation tests in southwestern Colombia.

Initially we carried out a pilot test to evaluate whether predators at these sites responded to the artificial models and whether they were attracted to a particular shape (elongated or spherical). For this test, 20 *P. dulce* separated by a distance of at least 30 m were used in each area (urban, suburban). Ten trees received sphere-shaped models (five on stems and five on leaves), and 10 received elongated larvae (five on stalk and five on leaf). Five models were placed on each tree. This test lasted for 30 days.

For the main predation test (PT), ten trees were selected and five artificial larvae were placed at heights of between 1.5 and 2 m, separated by at least 250 mm, and distributed according to substrate: five trees had larvae on their leaves and five on the stems. The test lasted for 30 days with two repetitions (October-November 2015 and January-February 2016) and each site was visited twice per week. At each visit, the larvae were moved to a different place on the same tree to avoid predator bias through learning, and all of the larvae with attack marks were replaced [13]. Initially, a total of 200 models were installed and the number increased as those with evidence of attack were replaced.

### Identification of potential predators

In order to identify the potential bird predators of the models, 10 observation and counting points, 15 m in diameter and 93 m apart, were establish at each site. Each site was visited and inspected for 15 min, between 7:00-10:00 h and 14:00-17:00 h. This activity was carried out once at each site during each experiment. The birds observed were identified to species following specialized keys [20] [21].

A single sampling of arthropods was carried out using three methods:

a. Pitfall traps [22] i.e. 10 plastic glasses containing 68 cc of alcohol installed at each site (one meter from the trees used in the predator experiment) and left in operation for 48 h;
b. Tuna bait [23–24]. Pieces of white paper (70 mm×220 mm) containing small quantities of tuna in oil were installed at observer height, one on each of 10 *P. dulce* trees at each sampling site. The bait was left from 14:00 h until 16:00 h at the end of which period the attracted insects were collected and placed in 95 % alcohol;
c. Two Malaise traps [25], i. e., a set of nets arranged in the shape of a field tent with an opening at the bottom, were placed at each site for an entire week. The arthropods were identified up to order, following taxonomic keys [26] while the predators were identified up to family and/or subfamily [27–28].

Recognizing predator marks. A guide to the marks on the models was produced (Fig 2). After finishing the field phase, 30 larvae models containing tuna in their interior as bait were displayed on five trees on the University campus. After two hours, the marks were compared with those reported by other colleagues [13]. In addition, an individual of *Polistes erytrocephalus*, a wasp species predominant in the area, was captured and placed near a model, pressing its sting against the plasticine in order to obtain bite and sting marks. All of the marks were photographed.

**Fig 2.**
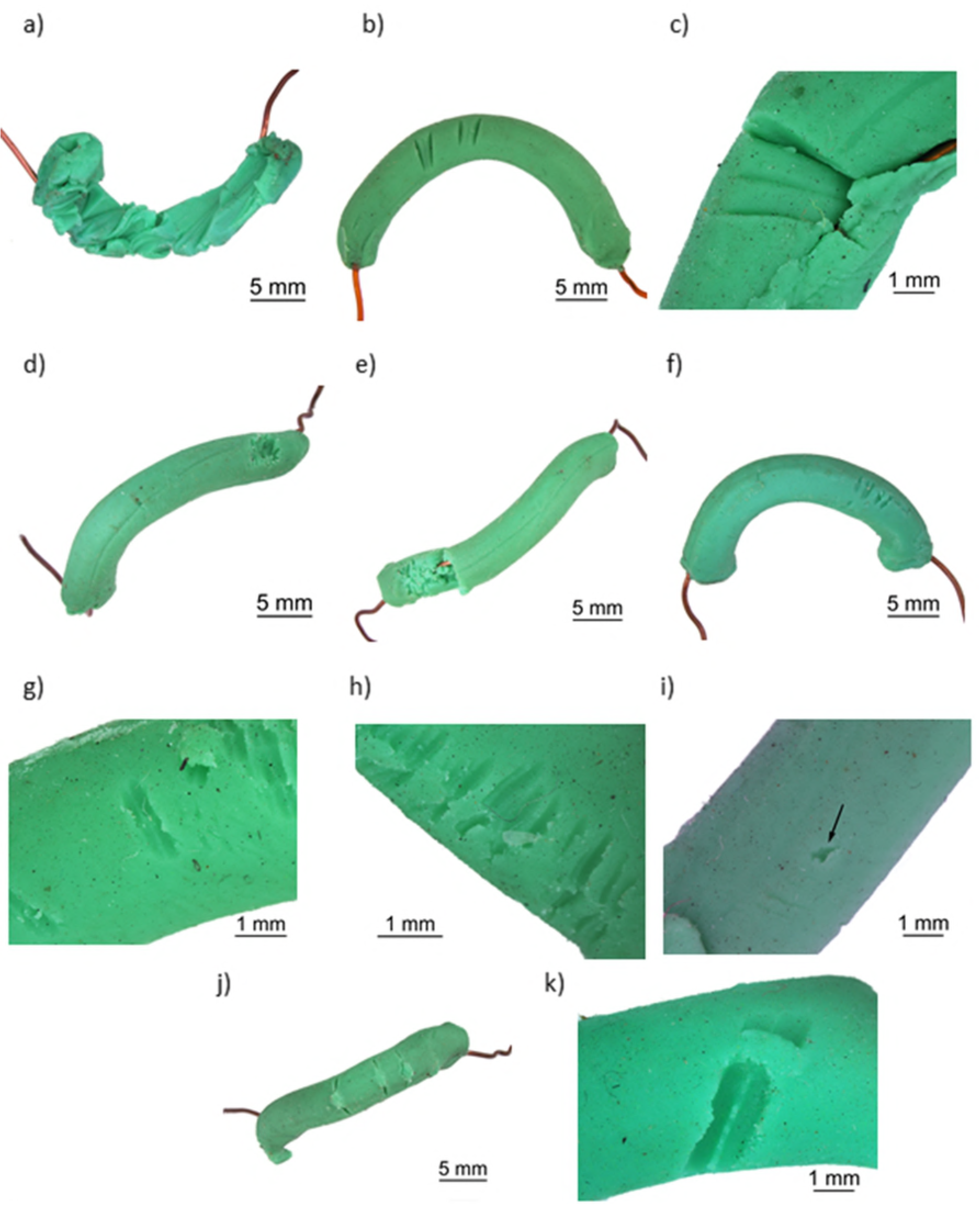
Predator marks registered on the lepidopteran larvae models. **a, b** and **c** bird beak; **d** and **e** ants; **f,g** and **h** wasp jaw; **i** wasp sting; **j** chewing arthropod and **k** mammal. Southwestern Colombia. Photo taken by Image Laboratory at the Graduate School of Biological Sciences, Universidad del Valle.

### Statistical analysis

Proportion of attacks by birds and arthropods on the artificial models: the two types of substrates and shapes of the model (elongated or spherical) were compared and analyzed using chi-squared tests [29–30]. A Generalized Linear Mixed Model (GLMM) was used, with predation as a binomial variable response and habitat (urban-suburban), substrate (stem-leaf) and season (dry-rainy) as explicative variables. We evaluated the possible relationship between predator abundance (birds-arthropods) and number of attacks carried out using linear regression. Species accumulation curves were used to evaluate the representativeness of the bird sampling, and the ACE estimator using the Estimates 9.1.0 Program [31]to measure expected richness. For arthropods, we evaluated the total number of individuals, to order level, captured at the sites evaluated and using the three methods described above. For Hymenoptera, the number of individuals per family and subfamily was indicated. R-Studio was used for all of the analyses [32]. Probabilities of less than or equal to 0.05 were considered significant.

## RESULTS

### Predation tests

In the pilot test, 23.48% of the models had predator marks and significant differences were detected between the proportion of attacks and the type of model and predator. Elongated models (n = 36, 62.07%) were attacked significantly more than spherical (n = 22; 37.93%, *χ*^2^ = 6.76, df = 1, P = 0.01) by predators. This difference is because arthropods did not attack any of the spherical models (elongated n = 8 or 22.2%; spherical = 0) while birds attacked both in similar numbers (elongated n = 28 or 77.8%; spherical = 22 or 100%; *χ*^2^ = 0.02). In the main test, of the 518 artificial models used, 125 (24.13%) showed evidence of predator attacks. No models were lost. Incidence of depredation was significantly greater in the urban area (63.20%) than in the suburban (36.80%). The leaf substrate showed a greater number of attacks than the stem (60% vs 40%). Season (rainy-dry) had no significant effect on incidence of predation nor was there an interaction effect among the variables used (Table 1). Birds were the most important predators (74.40%) followed by arthropods (24.80%) (Table 2). Only one mammal attack was registered (0.80%). There was a positive correlation between the abundance of birds (0.95) and the number of attacks on the models; the contrary was found for arthropods (0.47) (Fig 3).

**Fig 3.**
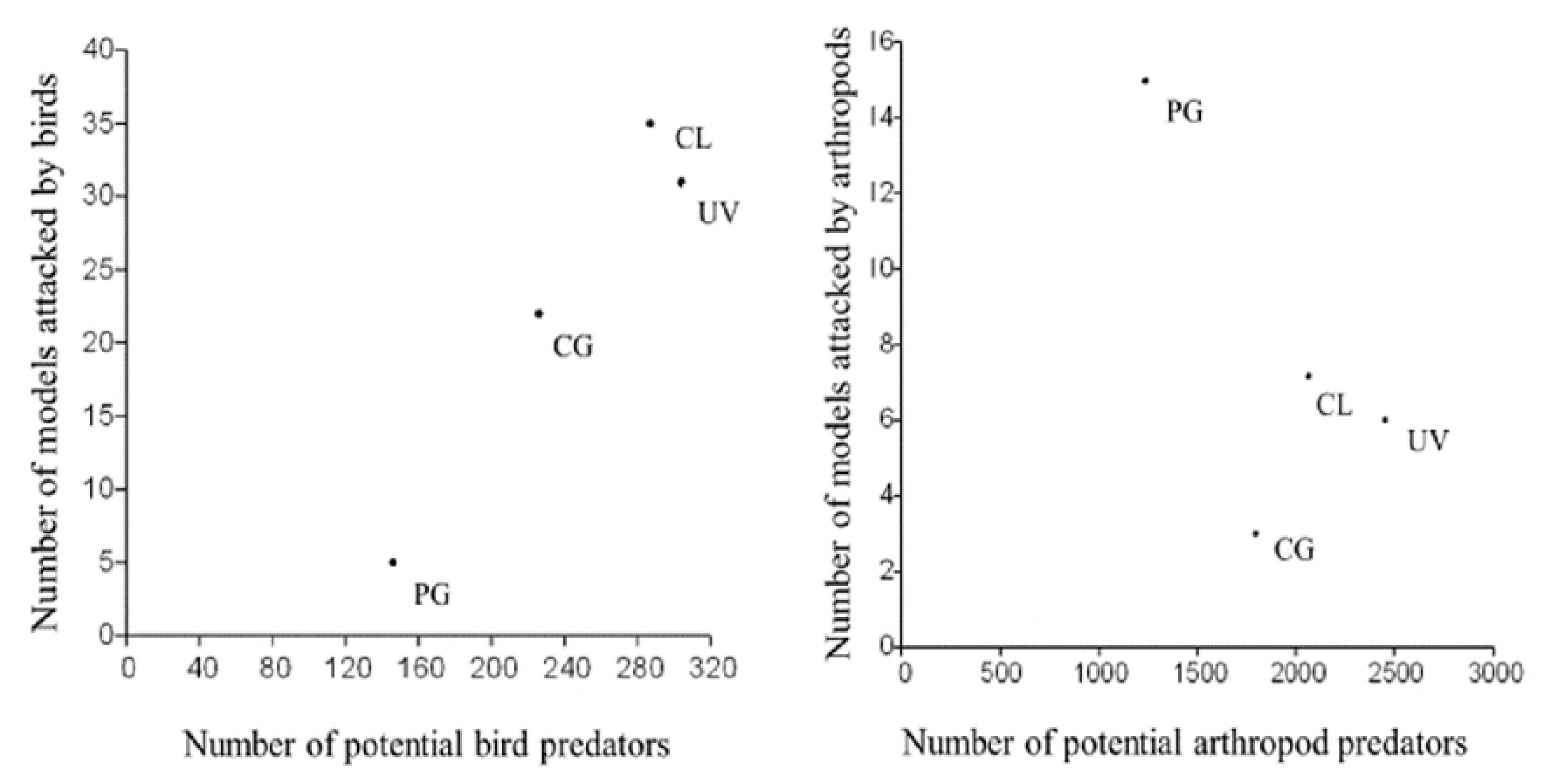
Correlation between predator’s abundance (birds-arthropods) and the attacked models. Correlation between bird abundance (a) and arthropods (b) with number of models attacked at each of the study sites (CG= Hacienda Cañas Gordas; PG= Parque de las Garzas; CL= Constructora Limonar; UV= Universidad del Valle) in Southwestern Colombia. Correlation coefficient: birds 0.95; arthropods 0.47.

**Table 1.** Generalized Linear Mixed Model (GLMM) where predation models of lepidopteran caterpillars were evaluated according to degree of urbanization (U: urban, SU: suburban), substrate (S: stem, L: leaf) and season (Ra: rainy, Dr: dry) in southwestern Colombia. Hab: habitat, Sit: site, Sub: substrate, Per: period. df = degrees of freedom; P = probability

| Variables | $\chi^2$ | df | P |
| --- | --- | --- | --- |
| **Period Ra-Dr** | 1.97 | 1 | 0.15 |
| **Habitat U-SU** | 5.65 | 1 | 0.01 |
| **Substrate S-L** | 3.82 | 1 | 0.05 |
| **Hab x Sit** | 0.82 | 2 | 0.66 |
| **Hsb x Sub** | 0.12 | 1 | 0.71 |
| **Per x Hab** | 2.51 | 1 | 0.11 |
| **Per x Sub** | 2.57 | 1 | 0.10 |
| **Per x Hab x Sub** | 2.70x10<sup>-3</sup> | 1 | 0.95 |

**Table 2.** Total number of attacks carried out by birds and arthropods on lepidopteran caterpillar models according to substrate in southwestern Colombia.

| **Birds** |  | **Arthropods** |  | **Total** |  |
| --- | --- | --- | --- | --- | --- |
| **Leaves** | **Stem** | **Leaves** | **Stem** | **Birds** | **Arthropods** |
| 55 | 38 | 20 | 11 | 93 | 31 |
| (59.14%) | (40.86%) | (64.52%) | (35.48%) |  |  |
| $\chi^2 = 6.22$, df = 1, P = 0.013 | | $\chi^2 = 5.23$, df = 1, P = 0.02 | | $\chi^2 = 62$, df = 1, P = 0.34x10<sup>-14</sup> | |

### Potential predators

#### Birds

A total of 74 bird species were found grouped in 30 families, Tyrannidae being the most representative. Species accumulation curves show 78.15% (suburban environment) and 75.13% (urban) efficiency in the avifauna sampled (S1 Fig). Of the total number of species, 42 were considered to be potential predators of lepidopteran larvae due to their foraging habits [20, 33–38] (S1 Table, S2 Fig). Abundance of this group of birds varied significantly with degree of urbanization, the urban area having the greater number of individuals (*χ*^2^ = 49.80; df = 1; p = 0.170×10^-11^).

#### Arthropods

Fifteen orders of arthropods were found, five of them potential predators: the most representative groups being Hymenoptera, Lepidoptera and Diptera (S2 Table). However, only Coleoptera and Hymenoptera were responsible for the arthropod attacks (24.80%) recorded in this study. Of Hymenoptera, the most abundant family was Formicidae (S3 Table) and, within it, Dolichoderinae and Myrmicinae were the most representative subfamilies (S4 Table). Wasps were responsible for 58.06% of attacks by arthropods (66.67% corresponding to bites and 33.33% to stings), 38.71% by ants and 3.23% by other chewing arthropods.

## DISCUSSION

### Degree of urbanization

The greatest incidence of predation on lepidopteran larvae models in the urban area when compared to the suburban area could be related to the increased abundance of birds associated with urbanization. In urban settings, the abundance of some bird species increases due to the absence or reduction of the predators that control them; in these environments, the survival of predators such as snakes, or birds of prey diminishes. Although the presence of other predators such as dogs and cats increase, the pressure exerted by them does not significantly affect the population density of urban birds [39]. The absence or reduction of their predators allow urban birds more time for foraging, and an increase in the number of individuals that exerts greater predatory pressure on insects [39–40]. Additionally, areas of urban vegetation favor increased abundance of birds since these patches act as connecting points between urban and suburban areas, and offer food resources for exploitation by birds [18–19]. Predatory incidence was not related to arthropod abundance since there was a weak correlation (0.47) between arthropod abundance and attacks. The response by arthropods may be greater in better conserved environments with greater vegetation complexity, such as forests [2, 11, 41]. In addition, a greater amount of light and vegetation complexity in open environments, such as the urban and suburban areas may facilitate localization of prey by birds [11,13,41]. The degree of habitat disturbance may have a significant effect on predation of insects by herbivores. In Panama, for example, it was found that the incidence of depredation on artificial larvae in forest clearings was significantly higher than that registered in closed forest [8]. This finding can be explained by forest clearings presenting high primary productivity, thus housing a larger number of herbivorous insects controlled by predators. In the Philippines, on the other hand, predatory incidence on herbivores was significantly greater in the rural area (59.4%) compared to areas with close canopy (46.1%) [11]. Equally, in Costa Rica, it was found that predation pressure on herbivores in open fields is double to that in forests [41].

In Denmark, at ground level, an increase in level of urbanization could reduce the predation incidence on caterpillars (43.98% forest, 30.77% suburban, and 25.25% urban) [2]. In this case, incidence of predation would be associated with level of habitat conservation since better-protected places would have a greater diversity of natural enemies. On the other hand, the results of our study suggest that an increased abundance of birds is responsible for higher predation rates [42–43]. In Papua Nueva Guinea while comparing the response of predators (birds-ants) along an altitudinal gradient it was found that predation rate was dependent on the number of species of insectivore birds and their abundances, just as on the number of individual ants on the trees sampled [42]. In Hungary, in oak forests, the structural heterogeneity of forests can increase the abundance of insectivorous birds and, in turn, increase the incidence of bird predation on artificial models of a lepidopteran species [43].

Although the phenological state of *P. dulce* was not evaluated in the present study, it is known to produce floral buds and flowers throughout the year and its phenophases are constant throughout the year [44]. Nevertheless, it is not known whether subtle differences exist in the phenological state of some trees according to level of habitat disturbance, and whether, in this case, trees with a greater supply of fruits would be found in the urban areas, thus stimulating both an increase in generalist bird visits as well as the probability of contact with larvae. It would be advisable for future studies to include this variable in order to determine whether the response is only due to the disturbance factor and not to the visit of generalist birds due to fruiting.

### Substrate

The significant differences found in type of substrate could be related to the foraging of the community of predators present in the habitats studied [45–46]. In the northern United States, 11 bird species that forage on forest foliage do so mainly on the leaves and not on the bark of trees [47]. It was also found that some species, such as the Vireonidae and Cardinalidae families, capture a large proportion of lepidopteran larvae in this substrate through a variable distance search pattern (close and far from the substrate). In the same way, solitary wasps hunt herbivorous insects such as lepidopteran larvae among the leaves [13, 27, 48]. In a Puerto Rican novel *Prosopis-Leucaena* woodland [49] birds prefer to forage for food on *P. dulce* since this tree houses a large quantity of arthropods associated with its foliage due to the high nitrogen content and small amount of hemicellulose in its leaves. At the Universidad del Valle, this plant is frequented by both resident and migratory birds, their main feeding activity there is the search for and consumption of insects (61.5%) compared to seed (29.7%), flower (4.4%), nectar (2.2%) and leaf (2.2%) consumption [18–19].

The type of substrate can influence level of detection of prey by predators if the prey is more conspicuous in a given substrate [10, 13, 42, 50–51]. In Papua New Guinea [13], a higher level of exposure of artificial larvae increased their incidence of depredation; those exposed on leaves were attacked more significantly than those hidden in rolled up leaves. In some environments, stem predation was found to be 1.9 times greater than on leaves [10]. However, this also depended on the type of predators that make up the community in the area [52]. The birds’ response detected in this study might be consequence of particular preferences of various groups of birds for food substrate [52]. The different shapes and sizes of peaks support this. In the case of the arthropods, it cannot be ruled out that the marks found by wasps and ants were due to a single species of each group. It was complex identify their marks at a finer level [52–53].

### Type of predator

The difference between the proportion of attacks by birds and (74.40%) arthropods (24.80%) highlights the importance of birds as controllers of herbivorous insects in urban settings where they are the main predators on larvae [51,54]. It is also thought that birds have a beneficial ecological function for plants by reducing the seasonal load of herbivores [42]. Additionally, abundance of birds increases with degree of urbanization and quickly responds to outbreaks of herbivory [40, 55]. In North America, for example [56], birds were responsible for 50-80% of the reduction of larvae that fed on cranberry (*Vaccinium myrtillus*).

The size of a model and its color can also influence incidence of predation since both determine the degree of prey detection and selection. Birds respond positively to increased prey size, having a strong impact at the end of the larval period; on the other hand, the effect is the opposite in arthropods since these mainly attack small individuals in their first instars [54, 57–58]. Regarding the color of the model used, green-colored preys are perceived as harmless by predators [41, 59]. Chemical signals released by plants when they are attacked such as volatile compounds, and other types of signals such as waste produced by herbivores, rather than visual signals can be more important for attracting arthropods that prey on herbivore invertebrates [42, 60–61]. Although chemical signals can also attract birds, the response may be stronger for arthropods [42, 62]. For instance, recruiting of ant species of the *Azteca* genus is greatest and more localized on *Cecropia obtusifolia* trees with artificially damaged leaves than on non-damaged ones [60]. Similarly, both birds and ants along the altitudinal gradient attacked larvae on leaves with simulated damage significantly more times than on leaves with no damage, but that those on damaged leaves were attacked more by ants [42]. In our study, the damage produced by manipulation of the leaves when placing the models is unlikely to have generated an important enough chemical response to strongly attract other arthropods predators known to be present in the study sites [53]. This indicates that predators were attracted to the models using visual rather than olfactory or chemical cues.

Wasps are known to be important controllers of herbivores. Some species of the Ichneumonidae and Braconidae families are parasitoids on lepidopteran larvae [57, 63] and other solitary species, such as those of the genus *Polistes* are visually oriented predators that feed on larvae [13, 27, 48, 64]. The majority of attacks on arthropods by wasps could indicate that these are more influenced by visual signals than ants when foraging. For instance, wasps presented greater percentages of rejection to the increased size of their prey compared to ants [64]. Wasps are also influenced by the color of their prey while the ants are not. This would indicate that size of prey could be a limiting factor for predators with a solitary life style but not for those with a level of social cooperation such as the ants. Color can be an indicator of palatability for wasps while it is a chemical signal for ants.

### Seasons

Although no significant differences were found between the two seasons (dry-rainy), this variable can influence predator and prey populations. During periods of primary productivity, such as the rainy season, herbivore insect populations experiment peaks of maximum abundance that coincide with the breeding season of some predators such as birds [8, 51, 65].

### Method

The use of artificial models is a good option for evaluating how trophic dynamics varies along a disturbance gradient [53]. However, our results should be interpreted with caution. The use of this method can underestimate, or overestimate, real predation pressure since the models do not possess many of the characteristics of real larvae (movement, chemical signals, anti-predation capacity, etc.) [13–14, 51, 53, 59]. Nevertheless, this does not mean that it is less informative than other methods since some studies did not find significant differences between real and artificial larvae [8, 66]. Moreover, in this study, as in other works [13], we found that predators can respond to models in the shape of their prey.

## CONCLUSIONS

Our results suggest that predation pressure on a prey organism can vary significantly according to level of disturbance, substrate where it is located, and type of predator. Level of disturbance increases the abundance of some predators such as birds and increases the possibility of larvae being preyed upon. The substrate where prey is found becomes a key aspect for their detection and will depend on their predators’ particular foraging behavior. Birds are the main controllers of herbivore larvae in urban environments due to a combination of their abundances and foraging behavior.

## Supporting information

**S1 Fig. Species accumulation curves obtained for the two environments evaluated.** a) Suburban, sampling efficiency 78,15 %. b) Urban, sampling efficiency 75,13 %, in southwestern Colombia.

(TIF)

**S2 Fig. Number of potential bird predators of lepidopteran larva by family in southwestern Colombia.** Bird families putative predators of the plasticine models (X-axis) and the number of species for each of these families (on the Y-axis).

(TIF)

**S1 Table**.List of bird species, trophic guilds (Fr: Frugivorous, In: Insectivores, Gr: Granivores, Cr: Carnivores, Nc: Nectarivorous, and Ao: Aquatic omnivore) and their abundances in the surveys carried out in each of the study sites (CG= Hacienda Cañas Gordas; PG= Parque de las Garzas; CL= Constructora Limonar; UV= Universidad del Valle) during this study.

(DOCX)

**S2 Table**. Groups or orders of arthropods captured at each of the sampling sites with the three methods used. *= Potential predator.

(DOCX)

**S3 Table**. Number of individuals captured by Hymenoptera family at each of the sites by the three methods used.

(DOCX)

**S4 Table**. Total individuals captured in each of the Formicidae subfamilies at each of the sites.

(DOCX)

## ACKNOWLEDGEMENTS

Thanks to A K. Tvardikova for methodological advice and to A W. Torres for statistical advice. Thanks to I. Castro, H. Álvarez-López and M. D. Heredia for their valuable recommendations and to Marcia Ditmann, I Castro, and N Bansal for help with English translation. Finally, thanks to Constructora Limonar, Administradora de Vallados, Fundación Cañasgordas, and to the Department and Graduate Biology School, Universidad del Valle for providing access to study sites.

## Author contributions

**Jefferson Cupitra-Rodríguez:** Conceptualization; Data analysis; Investigation; Methodology; Project administration; Resources; original draft; Validation; Visualization; Review & editing.

**Lorena Cruz-Bernaté:** Conceptualization; Supervision; Investigation; Validation; Review & editing.

**James Montoya-Lerma:** Supervision; Investigation; Validation; Review & editing.

